# PRC2 activates interferon-stimulated genes indirectly by repressing miRNAs in glioblastoma

**DOI:** 10.1101/681858

**Authors:** Haridha Shivram, Steven V. Le, Vishwanath R. Iyer

## Abstract

Polycomb repressive complex 2 (PRC2) is a chromatin binding complex that represses gene expression by methylating histone H3 at K27 to establish repressed chromatin domains. PRC2 can either regulate genes directly through the methyltransferase activity of its component EZH2 or indirectly by regulating other gene regulators. Gene expression analysis of glioblastoma (GBM) cells lacking EZH2 showed that PRC2 regulates hundreds of interferon-stimulated genes (ISGs). We found that PRC2 directly represses several ISGs and also indirectly activates a distinct set of ISGs. Assessment of EZH2 binding proximal to miRNAs showed that PRC2 directly represses miRNAs encoded in the chromosome 14 imprinted DLK1-DIO3 locus. We found that repression of this locus by PRC2 occurs in immortalized GBM-derived cell lines as well as in primary bulk tumors from GBM and anaplastic astrocytoma patients. Through repression of these miRNAs and several other miRNAs, PRC2 activates a set of ISGs that are targeted by these miRNAs. This PRC2-miRNA-ISG network is likely to be important in regulating gene expression programs in GBM.

## Introduction

PRC2 (polycomb repressive complex 2) is an epigenetic modifier complex that binds near transcription start sites and elsewhere in the genome and silences genes by methylating histone H3 at K27 through its histone methyltransferase component EZH2. (1). PRC2 can exert its regulatory impact on gene expression by forming regulatory networks involving direct interactions with its target genes and indirect mechanisms where it regulates other gene regulators which in turn regulate secondary PRC2 targets (2).

Several cancers including GBM, are characterized by upregulation of multiple components of the PRC2 complex including EZH2 (2-4). PRC2 could play an oncogenic role by silencing tumor-suppressive protein-coding genes or non-coding RNAs. miRNAs are a class of ∼22 nt short non-coding RNAs that play significant regulatory roles in disease and developmental pathways. Several miRNAs have been identified that play both oncogenic and tumor suppressive functions in multiple cancers including GBM (5,6).

The regulatory interplay between miRNAs and PRC2 components has been shown to play important roles in several disease and developmental pathways. In several contexts, miRNAs exert their regulatory impact by repressing PRC2 components (7,8). Similarly, PRC2 can also transcriptionally silence miRNAs to cause indirect activation of genes (2). Interestingly, miRNAs can also regulate gene expression by inhibiting or promoting PRC2 chromatin occupancy (9). Such regulatory crosstalk between PRC2 and miRNAs is underexplored in GBM.

Previously, we identified a regulatory network between PRC2 and miRNAs where PRC2 represses genes by activating miRNAs (10). Here, we investigated the role of PRC2-mediated repression of miRNAs in GBM-derived cell lines and bulk tumor samples. Through repression of miRNAs, PRC2 indirectly activates genes stimulated upon type I and type II interferon treatment (interferon-stimulated genes, ISGs). In contrast to previous findings in the mouse, we find that PRC2 directly represses miRNAs encoded in the DLK1-DIO3 locus in GBM (11). In addition to indirectly activating ISGs, we found that PRC2 also directly represses a distinct set of ISGs.

## Materials and methods

### Cell lines, reagents and treatments

T98G cells (ATCC-CRL-1690) were grown in EMEM with 10% FBS. Cells were maintained at 37°C and 5% CO2. For type I interferon treatment, cells were treated with universal type I IFN-α (PBL Assay Science, 11200) at 100 U/ml final concentration for 6 hr. For type II interferon treatment, cells were treated with IFN-γ (Millipore Sigma SRP3058) at 30 ng/ml final concentration for 48 hr.

### RNA-seq and data analysis

RNA from interferon treated T98G cells and primary tumor samples was extracted with TRIzol. Total RNA was then enriched for mRNA using NEXTFLEX Poly(A) Beads (Bioo 512980) as previously described (12). Enriched RNAs were then chemically fragmented and libraries were constructed using the NEBNext small RNA library preparation kit (NEB E7330) as previously described (13). Reads were aligned to the reference human genome version hg38 using Hisat2 and processed as previously described (10). Reads were then counted using featureCounts and differential expression was calculated using DESeq2 (14,15). miRNA libraries were constructed from primary tumor samples derived as previously described (12). Reads mapping to mature miRNAs (miRBase v21) were then counted using a custom script.

### Luciferase reporter assays

The 3’ UTR of target mRNAs spanning at least 500 bp around the predicted miRNA binding sites were cloned into the psi-CHECK2 vector (Promega, C8021) downstream of the Renilla luciferase gene (Supplementary Table 1). Mutagenesis was performed using the QuikChange II site directed Mutagenesis Kit (Agilent, 200523). WT and EZH2 −/− T98G cells were transfected with WT or mutant 3’ UTR reporters and harvested after 48 hours. Luciferase activity was then measured using the Promega dual luciferase kit and calculated as the ratio of luminescence from Renilla to Firefly. To determine 3’ UTR-dependent gene activation by EZH2, Renilla to Firefly ratios (R/F) of EZH2-activated genes were further normalized to R/F of PPIF, a gene not regulated by EZH2. To validate the interaction of specific miRNAs and EZH2-activated genes, cells were transfected with luciferase reporter vectors containing WT or mutant (miRNA binding site deletion) 3’ UTR sequences. Luciferase signals were then calculated as the ratio of Renilla to Firefly.

## Results

### PRC2 regulates interferon-stimulated genes

Through analysis of RNA-seq data we previously generated from EZH2 −/− and WT T98G cells, we identified 1834 EZH2-repressed and 1519 EZH2-activated genes (10). Gene ontology analysis of the EZH2-activated genes showed them to be enriched for genes involved in cellular response to interferon treatment (Fig. 1A). To verify this finding, we identified genes stimulated in response to type I interferon (IFN-α) and type II interferon (IFN-γ) and then checked for the number of interferon-stimulated genes (ISGs) that were regulated by EZH2. We considered genes that showed greater than 0.5 log_2_ fold induction in response to either IFN-α or IFN-γ as ISGs for further analysis. We found that 40% of the genes regulated by EZH2 were ISGs (*P* = 9.6e-89). 737 ISGs showed higher expression levels in response to loss of EZH2, suggesting that they were normally repressed by EZH2 (Fig. 1B). To check if these genes were directly repressed by EZH2, we determined the presence of EZH2 binding and H3K27me3 enrichment proximal to their transcription start sites by ChIP-seq (Fig. 1C, D). We found that EZH2 occupied the chromatin around 53% of the EZH2 repressed ISGs. Thus, consistent with previously published data, we found several ISGs to be repressed by EZH2 through its histone methyltransferase activity. However, we also found a distinct set of 630 ISGs to be downregulated upon loss of EZH2, suggesting they were activated by EZH2. Given that EZH2 primarily functions as a repressor we reasoned that the ISGs activated by EZH2 were activated indirectly. Since many ISGs are known to be regulated by miRNAs, we hypothesized that many EZH2-activated ISGs could be activated by EZH2 via repression of miRNAs.

**Figure 1.**
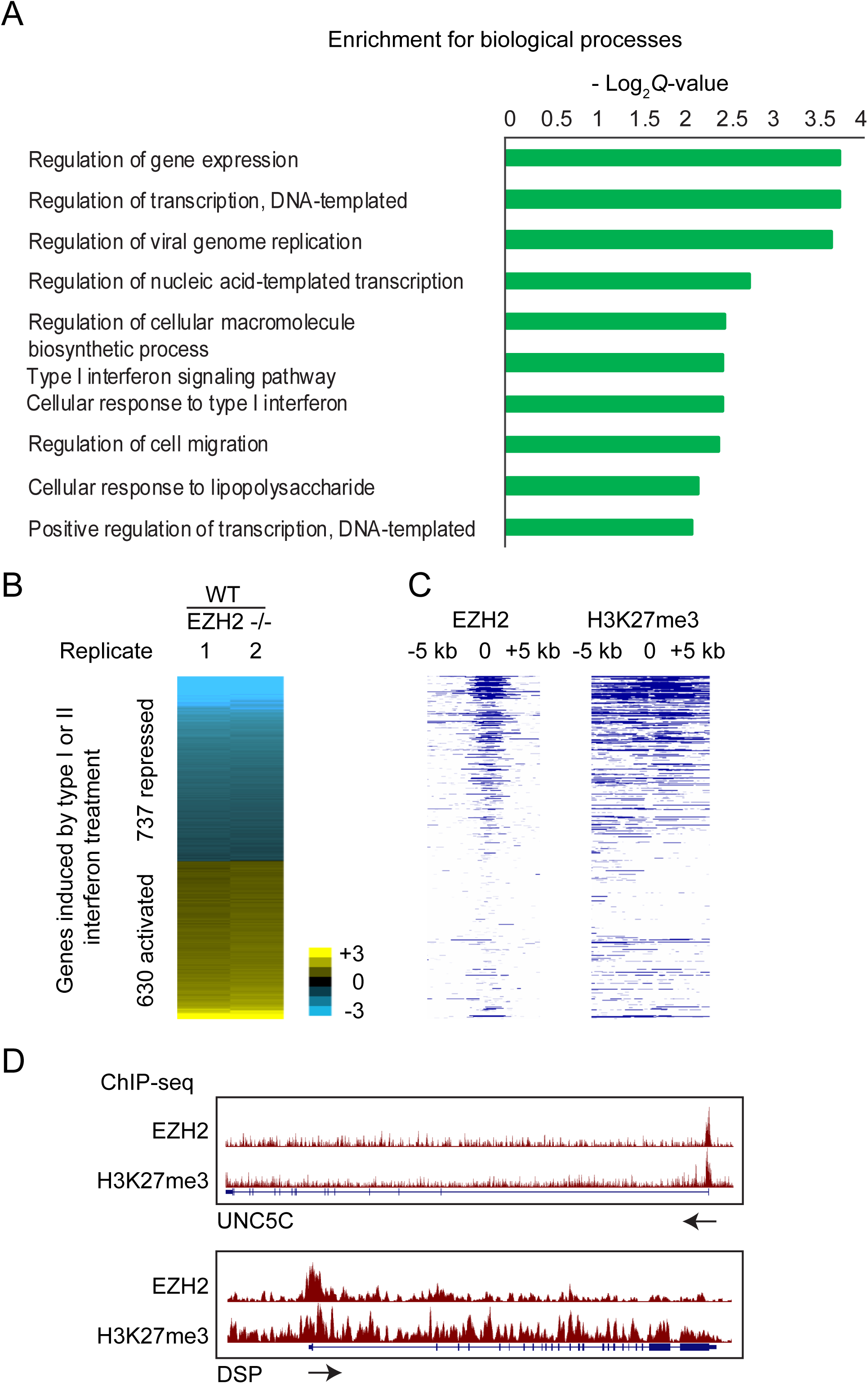
EZH2 represses ISGs directly and also activates them indirectly. Enrichment of biological processes among genes activated by EZH2, calculated by Enrichr. (B) Heat map showing differential expression of interferon-stimulated genes (ISGs) in response to EZH2 knockout. Genes are ranked in increasing order of log_2_ fold change (≤ −0.5 and ≥ 0.5 log_2_ fold change) in WT/EZH2 −/−. (C) Heat map showing enrichment scores (-log_10_ *Q*-value calculated using MACS2) for EZH2 and H3K27me3 for genes plotted in and ordered as in B. (D) Genome browser view of UNC5C and DSP showing tracks for EZH2 and H3K27me3 ChIP-seq normalized signal. Arrows indicate the direction of transcription.

### PRC2 directly represses miRNAs at the DLK1-DIO3 locus

Using the miRNA-seq data that we previously generated from EZH2 −/− and WT cells (10), we identified 84 miRNAs that were significantly repressed by EZH2. To determine if PRC2 silenced specific clusters of miRNAs, we calculated the frequency of EZH2-repressed miRNAs per chromosome. Chromosome 14 stood out in showing the highest proportion of EZH2-repressed miRNAs (Fig. 2A). 52 of the EZH2-repressed miRNAs mapped to the DLK1-DIO3 imprinted locus on chromosome 14. 67 of the 68 mature miRNAs expressed from the DLK1-DIO3 imprinted locus showed a trend towards repression by EZH2 (Fig. 2B). We found enrichment for EZH2 and H3K27me3 at this locus in T98G GBM cells (Fig. 2C), indicating that PRC2 directly and coordinately represses this cluster of miRNAs by methylating H3K27 across this locus.

**Figure 2.**
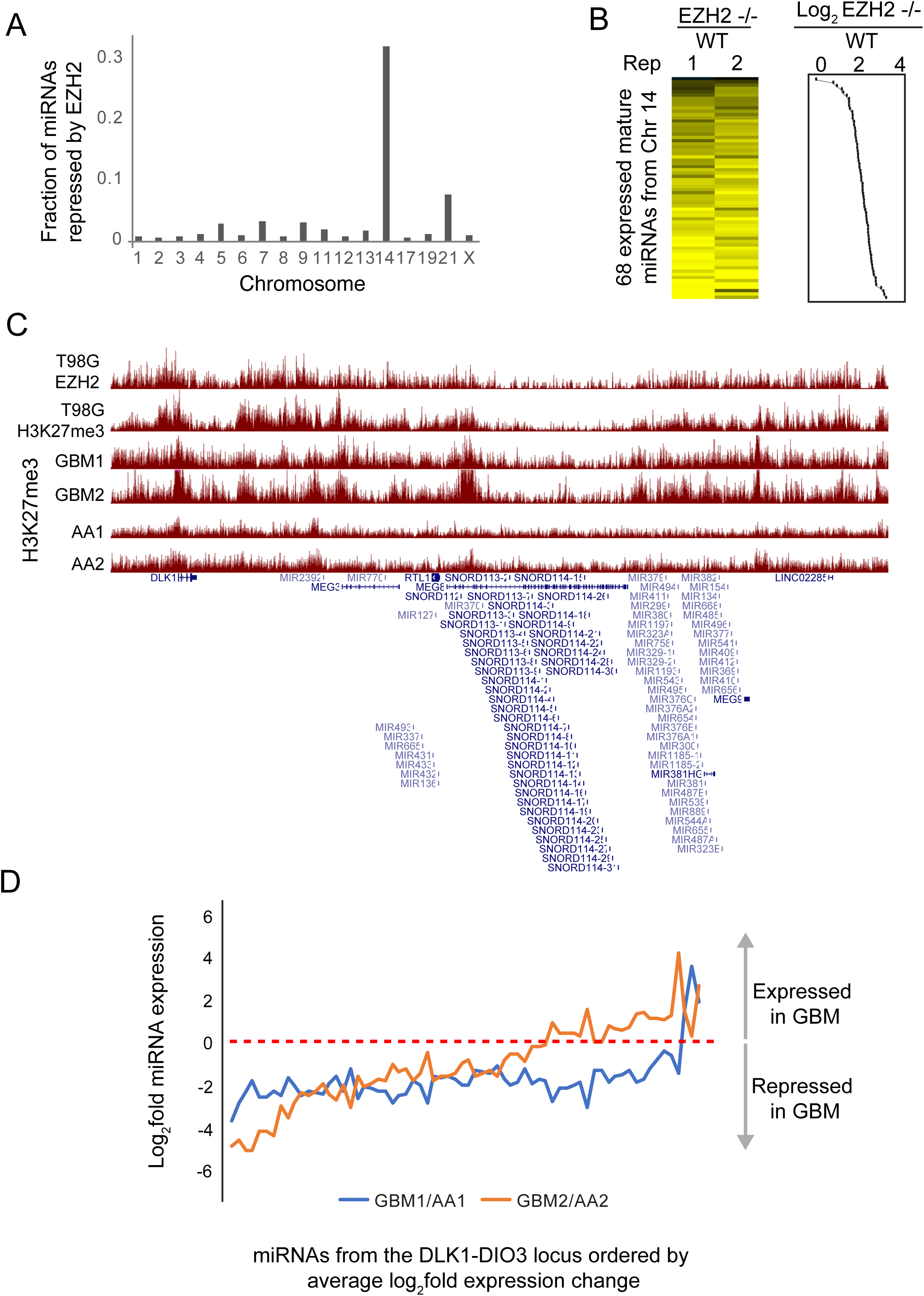
PRC2 directly represses the DLK1-DIO3 miRNA cluster. A) Fraction of miRNAs repressed by EZH2 per chromosome calculated as the ratio of the number of EZH2-repressed miRNAs and the number of miRNAs encoded per chromosome. (B) Heat map showing fold change of DLK1-DIO3 locus encoded miRNAs between EZH2 −/− and WT T98G cells. Line plot showing average log_2_ fold change is plotted to the right of the heat map. (C) Genome browser view showing EZH2 and H3K27me3 enrichment at the DLK1-DIO3 locus in the T98G GBM cell line and in primary tumors from GBM (GBM1, GBM2) and anaplastic astrocytoma (AA, AA2) patients. (D) Line plot showing fold change between normalized miRNA read counts for two sets of samples – GBM1 and GBM2, relative to AA1 and AA2, respectively. The X-axis shows miRNAs ordered by their average fold change across the two comparisons.

To determine if PRC2 also regulated the DLK1-DIO3 locus-encoded miRNAs in primary brain tumors, we analyzed H3K27me3 ChIP-seq data that we had previously generated in primary tumors (12). We found H3K27me3 to be differentially enriched at this locus across multiple GBM (GBM1 and GBM2) and anaplastic astrocytoma (AA1 and AA2) tumor samples (Fig. 2C). To test if low enrichment of H3K27me3 could lead to upregulation of miRNAs encoded in this locus, we performed miRNA-seq for GBM1, GBM2, AA1 and AA2. Relative to AA1 and AA2, both GBM1 and GBM2 showed lower expression for 94% and 67% of the DLK1-DIO3 locus-encoded miRNAs, respectively (Fig. 2D).

### PRC2 activates genes through repression of miRNAs

To determine the role of PRC2-mediated repression of miRNAs encoded in the DLK1-DIO3 locus, we identified genes that were activated by EZH2 via repression of miRNAs. The canonical targeting mechanism of miRNAs involves miRNA-mRNA pairing at the 3’ UTR of the target mRNAs. If EZH2 activated genes by inhibiting expression of miRNAs that target them, we would expect the activation by PRC2 to be dependent on the 3’ UTR. To test this, we performed luciferase reporter assays where we cloned the 3’ UTR region of EZH2-activated genes and compared luciferase expression in EZH2 −/− and WT cells. If the activation by EZH2 was 3’ UTR-dependent, we would expect the luciferase signal to be lower in EZH2 −/− cells due to higher expression of EZH2-repressed miRNAs compared to WT cells (Fig. 3A). Eight genes, including 7 ISGs, out of the 17 EZH2-activated genes that we tested showed a dependence on their 3’ UTR for EZH2 activation. This data suggests that these genes are under greater repression by miRNAs in EZH2 −/− cells compared to WT cells (Fig. 3B).

**Figure 3.**
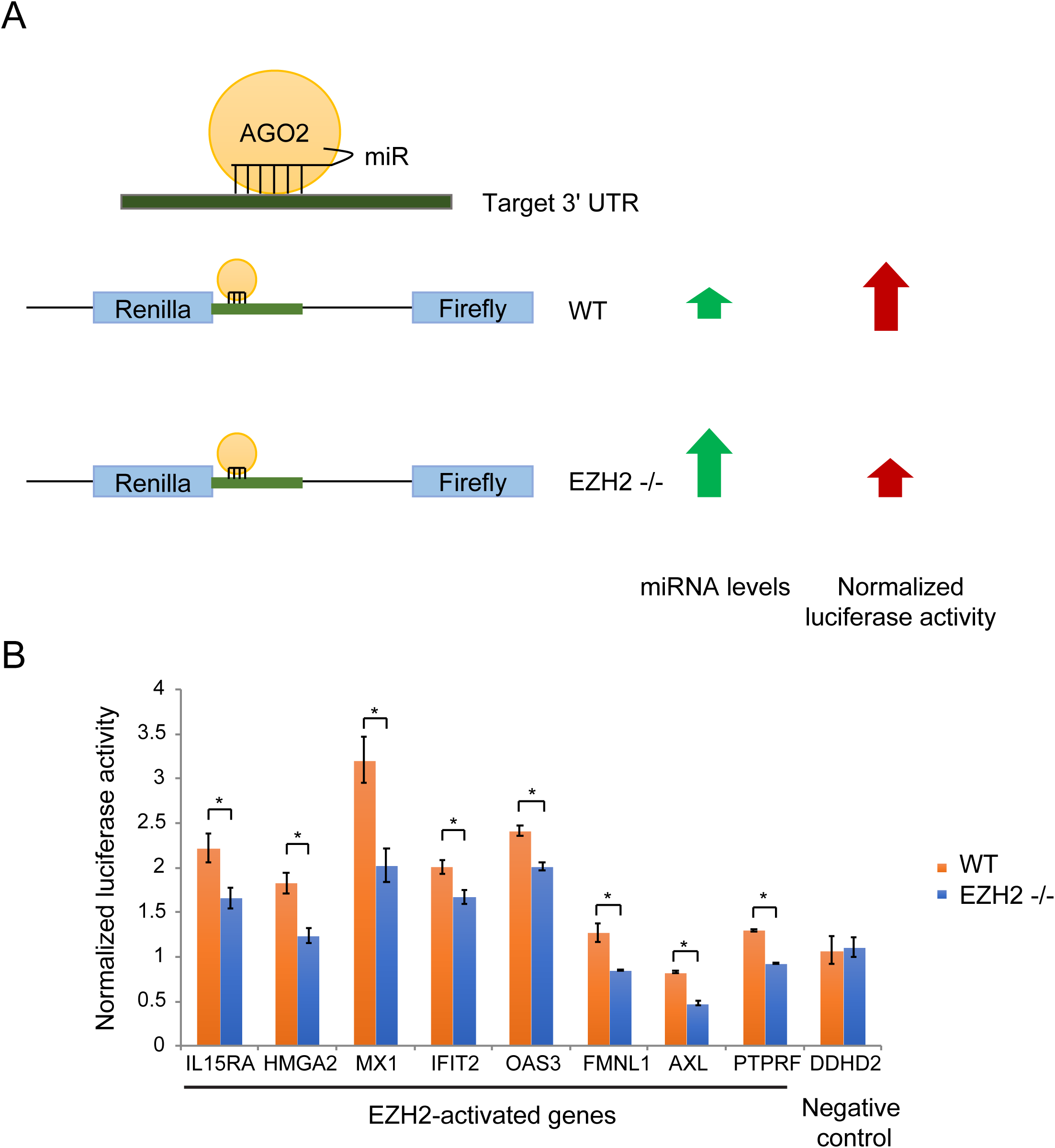
Genes activated by EZH2 in a 3’ UTR-dependent manner. (A) Schematic of the luciferase reporter assay to identify genes regulated by EZH2 in a 3’ UTR-dependent manner. (B) Column plots showing normalized luciferase signal from WT and EZH2 −/− cells overexpressing luciferase reporter constructs containing 3’ UTR regions of genes indicated on the X-axis.

We then verified the interaction between three EZH2-activated mRNA and EZH2-repressed miRNA pairs - HMGA2: miR-204-5p, MX1: miR-204-5p and IL15RA: miR-370-3p, in two ways. First, we performed luciferase reporter assays where we transfected EZH2 −/− cells with luciferase constructs containing WT or mutant 3’ UTRs that lacked the miRNA binding site. Deletion of miRNA-binding sites led to a rescue in luciferase expression (Fig. 4A). Second, we tested if the miRNA binding site deletion abolished the effect of the loss of miRNAs. We utilized cells stably overexpressing VP55, a vaccinia virus-derived poly(A) polymerase, to globally deplete miRNAs (16). IL15RA and MX1 WT 3’ UTR showed a rescue in luciferase signal upon miRNA depletion but the constructs with the mutant 3’ UTR did not (Fig. 4B). Taken together, this data shows that IL15RA is directly targeted by miR-370-3p and MX1 and HMGA2 are directly targeted by miR-204-5p. Of these miRNAs, miR-370-3p maps to the DLK1-DIO3 imprinted locus on chromosome 14.

**Figure 4.**
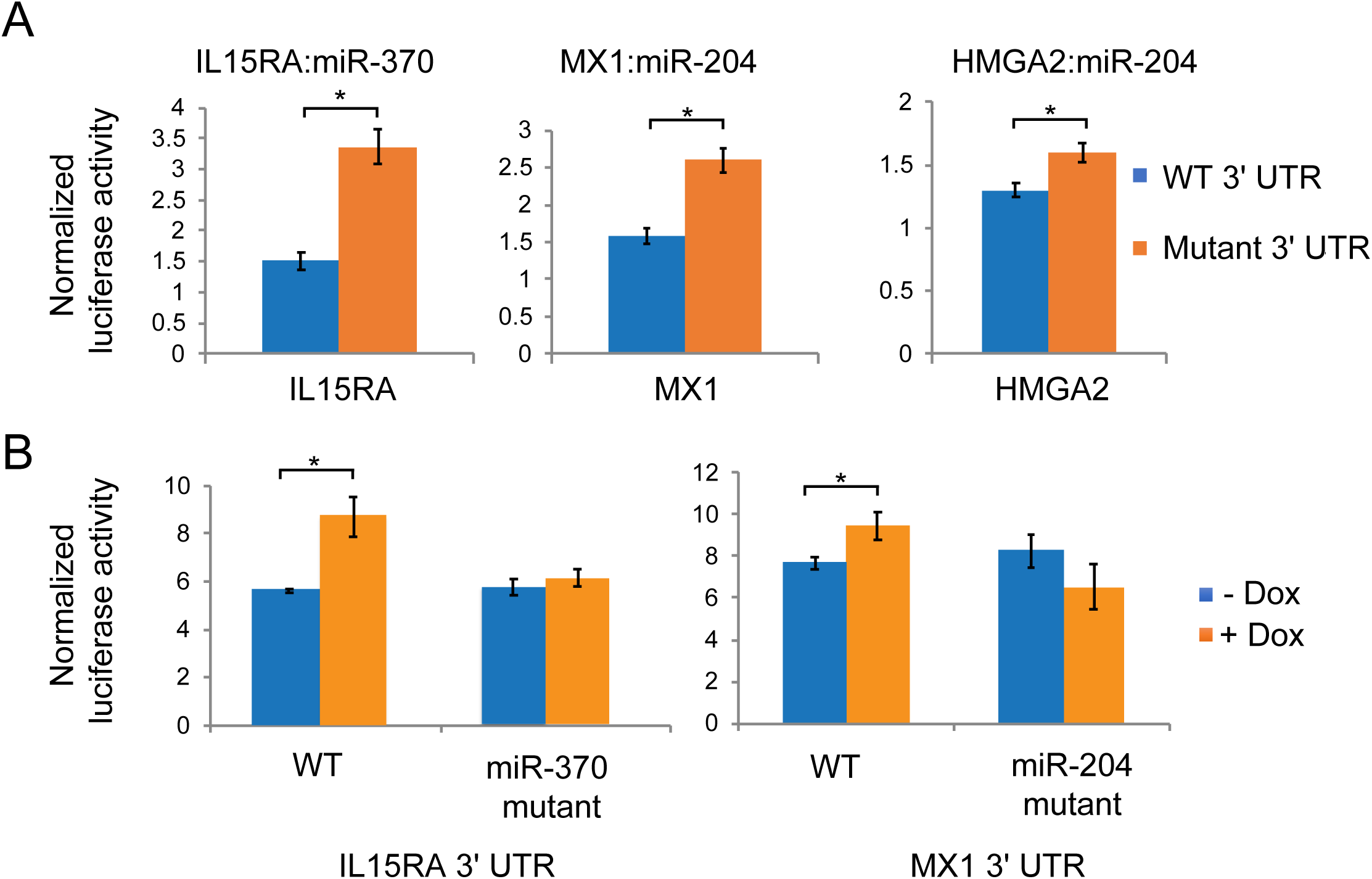
Direct regulation of EZH2-activated genes by EZH2-repressed miRNAs. (A) Column plots showing normalized luciferase signal from EZH2 −/− cells expressing luciferase reporter constructs containing WT and mutant 3’ UTR of IL15RA, MX1 and HMGA2 lacking the binding site for their targeting miRNAs. (B) Plotted as in (A) for WT and mutant 3’ UTR-transfected EZH2 −/− cells depleted of miRNAs by VP55 overexpression (+ Dox) or controls (-Dox).

To identify additional genes that could be indirectly activated by EZH2 via repression of miRNAs, we analyzed RNA-seq datasets from EZH2 −/− and WT cells overexpressing VP55 (10). We calculated the response to miRNA loss as the ratio of gene expression in cells depleted of miRNAs (+ Dox) to their expression in control cells (no Dox). If EZH2 activated genes by repressing miRNAs, we would expect those genes to be derepressed to a greater extent in response to loss of miRNAs in EZH2 −/− cells than in WT cells. With this approach we identified 213 EZH2-activated genes including 113 ISGs, that showed greater derepression, or higher induction in expression in EZH2 −/− compared to WT cells. This suggests that these genes are more strongly repressed by miRNAs in EZH2 −/− cells than in WT cells, likely due to higher expression of miRNAs in EZH2 −/− cells (Fig. 5).

**Figure 5.**
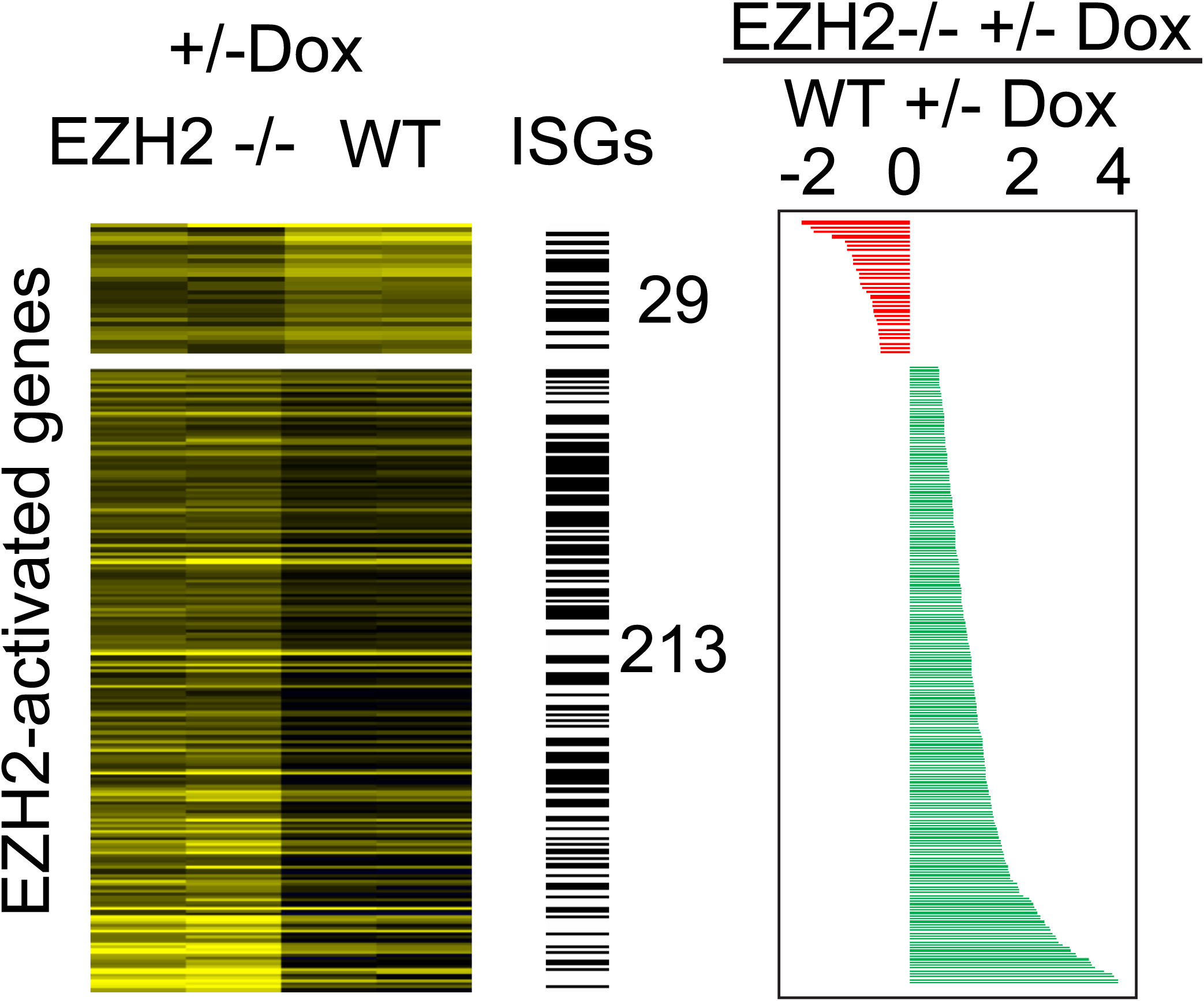
Identification of genes activated by EZH2 via repression of miRNAs. Heat map showing response to loss of miRNAs calculated as the ratio of normalized read counts in VP55 overexpressing (+ Dox) and control cells (-Dox), in EZH2 −/− and WT T98G cells. ISGs are indicated by black bars. The plot to the right shows the ratio of the response to miRNA loss in EZH2 −/− to WT cells. Genes indicated in red columns show lower response and genes indicated by green columns show higher response to miRNA loss in EZH2 −/− compared to WT cells.

## Discussion

The role of PRC2 in gene regulation has largely been focused on its direct repression activity (4,17,18). However, PRC2 could also activate genes indirectly by repressing other repressors. We found that EZH2 directly represses a set of interferon-stimulated genes (ISGs) and also activates a distinct set of ISGs through repression of miRNAs. We first identified miRNAs that were repressed by EZH2 and then determined the EZH2-activated genes that these miRNAs directly regulate. For several other EZH2-activated genes we determined regulation by miRNAs using a new approach that involves global degradation of miRNAs by overexpression of a vaccinia virus-derived poly(A) polymerase, VP55 (16).

DLK1-DIO3 locus is a maternally imprinted locus that when deregulated can lead to several developmental disorders including cancer (11,19). This locus, on chromosome 14 in the human genome and chromosome 12 in the mouse, encodes several protein-coding and non-coding genes including snoRNAs and miRNAs. Nine of the miRNAs encoded in this locus have been shown to be downregulated in GBM tumors compared to normal brain samples (19). We found that 67 of the 68 miRNAs encoded in this region to be directly silenced by PRC2 in primary tumors, consistent with the fact that EZH2 is frequently activated in GBM. This is in contrast to mouse embryonic stem cells where PRC2 was shown to activate this locus by blocking DNA methylation (11). To further investigate the function of the PRC2-mediated repression of miRNAs encoded by the DLK1-DIO3 locus, we focused on the genes that were activated by EZH2. EZH2-activated genes showed enrichment for type I and II interferon responsive genes. We identified genes induced upon type I and II interferon treatment in GBM cells and found that 40% of the EZH2-activated genes were induced by interferon treatment. Consistent with previously published data, a significant fraction (∼40%) of the EZH2-repressed genes also showed induction on interferon treatment (20). This suggests that EZH2 represses ISGs directly and also activates them indirectly by repressing miRNAs that target them. Thus, the two key findings of this study are first, that PRC2 directly represses miRNAs encoded by the DLK1-DIO3 locus in humans, and second, that through repression of miRNAs, PRC2 activates ISGs in GBM.

Polycomb proteins and miRNAs have independently been implicated in the regulation of interferon genes (16,20,21). miRNAs have also been shown to activate expression of interferon pathway genes via repression of polycomb proteins (21). Conversely, we show that PRC2 activates ISGs by repressing miRNAs that target them, suggesting that PRC2 can both activate as well as repress expression of ISGs. A similar dual regulatory impact on interferon pathway genes was previously observed for SETD2 histone methyltransferase (22). It is possible that, like SETD2, PRC2 also functions in the fine-tuning of interferon response. Loss of PRC2’s histone methyltransferase activity leading to differential regulation of ISGs was previously shown to suppress viral infection (23). How PRC2-mediated regulation of interferon genes relates to antiviral activity will require further investigation.

## Data Access

Primary sequencing data generated in this study is available at NCBI’s GEO database (https://www.ncbi.nlm.nih.gov/geo/query/acc.cgi?acc=GSE133211).

## Acknowledgments

We thank Anna Battenhouse for assistance with aligning sequencing data and the Genomic Sequencing and Analysis Facility at UT Austin and the MD Anderson Cancer Center-Science Park NGS Facility for Illumina sequencing. The Science Park NGS Facility was supported by CPRIT Core Facility Support Grant RP120348. We also thank the Texas Advanced Computing Center (TACC) at UT Austin for the use of computational facilities.

---

**Supplementary Table 1.**
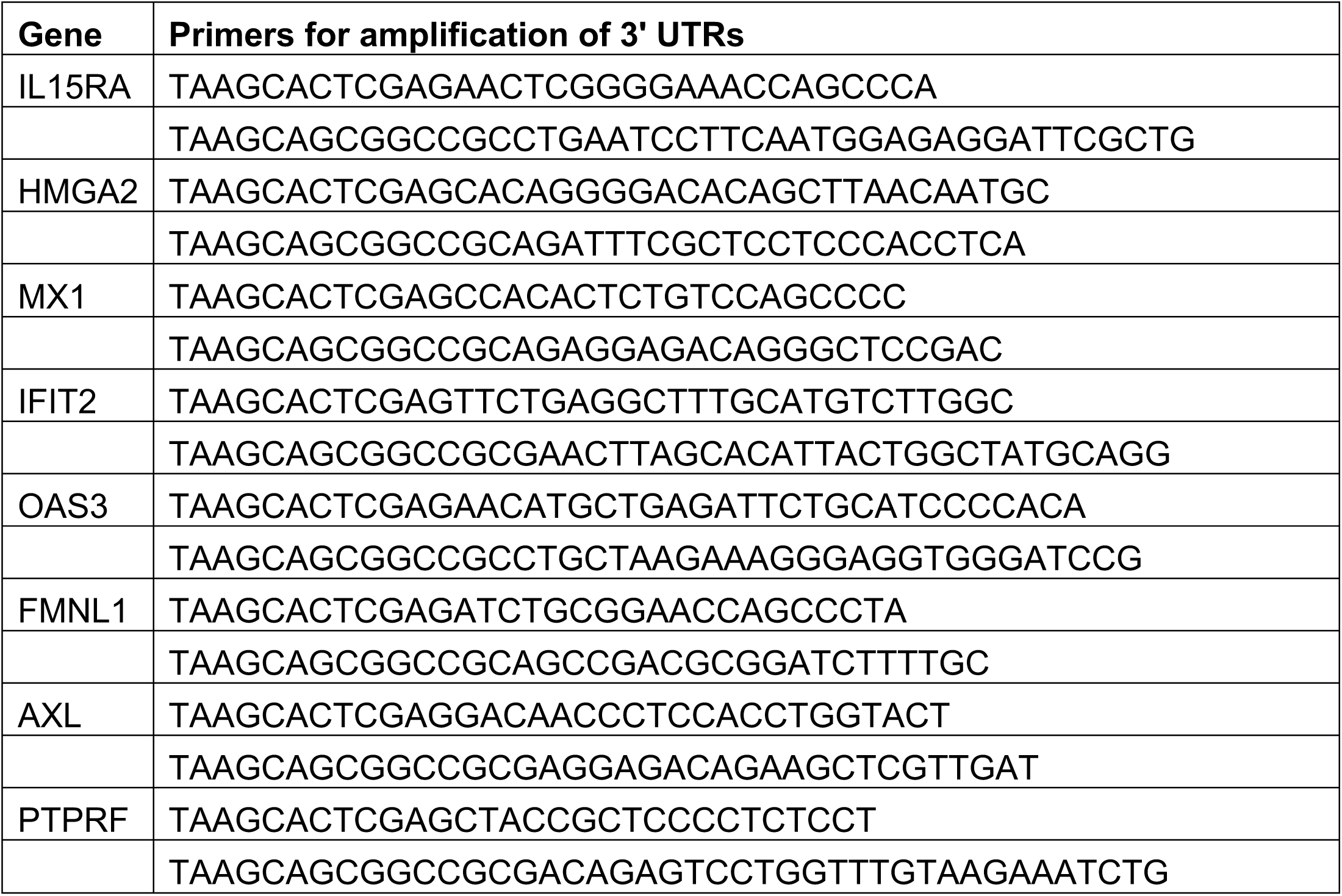
Primers for the amplification of 3’ UTRs of EZH2-activated genes

